# Different kinetics of humoral response against individual antigens in the ovine model of *Toxoplasma gondii* infection and its pathological association

**DOI:** 10.1101/2025.09.15.676282

**Authors:** Paula Formigo, Ignacio Gual, Germán J. Cantón, Luisa F. Mendoza Morales, Emiliano Sosa, Agustina Soto Cabrera, Valeria Scioli, Claudia Morsella, Valeria A. Sander, Dadin P. Moore, Sergio O Angel, Lucía M. Campero, Marina Clemente

## Abstract

**Background:** *Toxoplasma gondii* is a globally distributed protozoan parasite with major implications for human and animal health. In sheep, infection can cause reproductive failure and represents an important source of zoonotic transmission through contaminated meat. While cell-mediated immunity is the main protective mechanism, the contribution of humoral responses, particularly against individual recombinant antigens (rAGs), remains poorly characterized in ovine toxoplasmosis.

**Methods:** We analyzed the humoral response of lambs experimentally infected with two reference strains of *T. gondii* (RH and Me49) using different inoculation doses. Recombinant antigens rGra4Gra7, rGra8, rRop2, rCST9, and rBCLA were tested by IgG-ELISA to evaluate their kinetics during infection. Histopathological analyses of brain tissues were performed to assess lesion severity and cyst presence. Additionally, sera from naturally infected sheep were evaluated to test diagnostic performance.

**Results:** An early IgG response against rGra4Gra7 and rGra8 was detected, consistent with markers of acute infections. Anti-rBCLA, rRop2, and rCST9 IgG responses were associated with the chronic phase and correlated with cyst burden. Notably, stronger anti-rGra4Gra7 IgG responses tended to associate with lower lesion scores, suggesting a potential protective role. In naturally infected sheep, IgG-ELISA against rGra4Gra7 and rGra8 demonstrated the highest diagnostic performance, outperforming total lysate antigen (TLA)-based assays.

**Conclusion:** Our findings highlight distinct kinetics of specific antibody responses against selected *T. gondii* antigens in sheep, suggesting their value as markers of infection stage and pathology. rGra4Gra7 and rGra8 appear particularly promising both as diagnostic candidates and for exploring mechanisms of humoral modulation in ovine toxoplasmosis.

## 1 Introduction

Ovine toxoplasmosis is a disease caused by the obligate intracellular protozoan *Toxoplasma gondii*. In small ruminants, acute infection typically presents with a short episode of fever, apathy, anorexia, diarrhea, and coughing (1–3). In pregnant ewes, tachyzoites invade and replicate within the tissues of the feto-maternal junction (4), leading to severe reproductive outcomes such as abortion, fetal mummification, maceration, stillbirth, premature delivery, or the birth of weak lambs with low survival rates (5). The economic losses associated with toxoplasmosis in sheep flocks are substantial (6,7).

The course of *T. gondii* infection in intermediate hosts as small ruminants, is characterized by two distinct stages: an acute phase during the initial weeks post-infection, followed by a chronic phase marked by the formation of tissue cysts. In the murine model, chronic infection starts after the first week postinfection, but becomes established after approximately 20 days (8). In sheep, experimental infection with *T. gondii* oocysts results in the appearance of cysts in brain and muscle tissues about six weeks post-infection (9). These tissue cysts represent a major source of zoonotic transmission to humans, primarily through the consumption of undercooked meat or meat products (10).

Foodborne diseases (FBDs) are among the leading causes of morbidity and mortality worldwide, affecting nearly one-third of the global population annually (11). Although parasitic infections have historically received less attention than bacterial, viral, or chemical foodborne threats, they contribute substantially to the global burden of disease, as defined by the World Health Organization (WHO). Both the Food and Agriculture Organization (FAO) and WHO have highlighted the critical importance of foodborne parasitic diseases at the human–animal interface. This issue is highly relevant not only for public health, due to the spread and consequences of foodborne infections, but also for livestock production systems, where outbreaks can result in significant economic losses (12). Within this framework, the One Health approach has gained increasing importance by emphasizing the interconnectedness of human, animal, and environmental health in the prevention of zoonotic diseases. Among these, toxoplasmosis stands out as a major concern (7).

The host immune response against *T. gondii* is primarily mediated by a strong Th1-type response, characterized by the production of IFN-γ, which is essential for controlling parasite dissemination and disease severity (3,13,14). Nevertheless, some evidence indicates that B cells and the humoral response also contribute to protective immunity, albeit to a lesser extent. For instance, B-cell-deficient mice vaccinated with attenuated *T. gondii* strains exhibited greater susceptibility to challenge with virulent parasites compared to wild-type controls (15). Furthermore, passive transfer of IgG from mice immunized with ROP18 from *T. gondii* or from rats immunized with dense granule proteins conferred partial protection in experimental infections (16,17).

The humoral response elicited during *T. gondii* infection is highly consistent and easily measurable. Commonly applied serological methods include the Indirect Fluorescent Antibody Test (IFAT), Enzyme-Linked Immunosorbent Assay (ELISA), and various agglutination-based tests (18). Single-antigen approaches can provide advantages for diagnostic purposes as well as for monitoring infection dynamics and identifying clinical biomarkers (19). Antigens derived from rhoptries (ROPs), dense granules (GRAs), and surface proteins generally show the strongest diagnostic performance, while bradyzoite-specific antigens such as BAG1 have also been investigated (20,21).

Previously, we have characterized the antigenic value of multiple recombinant antigens (rAGs). Among them, rGra4, rGra7, rGra8, and the rGra4Gra7 chimera proved to be reliable markers of recent *T. gondii*-infection in humans and of acute stages in experimentally infected mice, outperforming rRop2 and the microneme antigen rMic1 (21,22). More recently, rCST9, a cyst wall/dense granule protein, was identified as an acute-phase marker, though with lower sensitivity (23). Notably, the rGra4Gra7 chimera also demonstrated value for monitoring acute infection in sheep experimentally infected with the RH strain (24). Additionally, rBCLA has been identified as a marker of chronic infection in both human sera and experimental mouse models (25), with elevated anti-BCLA IgG levels correlating with cyst-associated pathologies, particularly in immunosuppressed patients and those with ocular toxoplasmosis.

In this study, we evaluated the humoral response to rGra4-Gra7, rGra8, rRop2, rCST9, and rBCLA after experimental ovine toxoplasmosis with different *T. gondii* strains, as well as the relationship between pathological findings and antibody kinetics against the recombinant antigens (rAGs). In addition, serum samples from naturally infected and uninfected lambs were included to validate the experimental findings under field conditions.

## 2 Materials and methods

### 2.1 Parasite strains and culture conditions

RH RHΔhxgprt (26) and Me49 (23) *T. gondii* strains were grown in standard tachyzoite conditions *in vitro*: hTERT (ATCC® CRL-4001, USA) monolayers were infected with tachyzoites and incubated with Dulbecco’s modified Eagle medium (DMEM, Invitrogen) supplemented with 1% fetal bovine serum (FBS, Internegocios S.A., Argentina) and penicillin (10,000 units/ml)-streptomycin (10 mg/ml) solution (Gibco, Argentina) at 37°C and 5% CO2.

### 2.2 Experimental infection of lambs

Eighteen *T. gondii* seronegative, three-month-old Texel lambs from a flock at the National Institute of Agricultural Technology (INTA), Balcarce, Argentina, were selected for the experiment. Animal procedures were approved by the Institutional Committee for the Care and Use of Experimental Animals (CICUAE, INTA CeRBAS, protocol number 244/2022). The animals were housed in closed pens and provided with commercial alfalfa pellets and water *ad libitum*. Before infection, all lambs were bled, and their seronegative status for *T. gondii* and *Neospora caninum* was confirmed by indirect fluorescence antibody test (IFAT) testing (cut-off titer ≥1:50) (27). All animals were in good body condition, and their health was monitored throughout the study.

The lambs were randomly allocated into four experimental groups (n = 3 per group) and inoculated subcutaneously with different doses and strains of *T. gondii* tachyzoites: 5 × 10^6^ or 5 × 10^7^ of either the RH or ME49 strain. Before inoculation, tachyzoites were purified using a 3 μm polycarbonate filter and counted in a Neubauer chamber. Rectal temperature was recorded daily during the first 10 days post-infection. Two additional control groups were included for temperature analysis: three naïve lambs and three lambs subcutaneously injected with PBS.

Blood samples were collected on days 0, 7, 15, 30, and 60 post-inoculation. On day 60, all animals were euthanized in accordance with CICUAE guidelines. Euthanasia was performed by stunning with a captive-bolt pistol, immediately followed by exsanguination via the jugular vein. Necropsies were performed immediately after death was confirmed.

### 2.3 Recombinant antigens

The origin, expression, and purification procedures for rGra4Gra7, rGra8, rRop2, and rCST9 have been previously described (22,23). To generate rBCLA, a fragment of the corresponding gene was synthesized by GenScript (Piscataway, NJ, USA) and cloned into the pET28+ vector, as previously reported (25). Recombinant proteins were expressed in *Escherichia coli* cultures induced with IPTG and purified under denaturing conditions using Ni-NTA affinity chromatography (Ni^2+^-nitrilotriacetic acid resin, Qiagen, Valencia, CA, USA), following the manufacturer’s instructions. Protein purity was assessed by SDS-PAGE, and concentrations were determined by the Bradford assay.

### 2.4 Serum samples

To validate the experimental findings, serum samples were obtained from naturally infected and uninfected lambs belonging to a flock at Instituto Nacional de Tecnología Agropecuaria (INTA), Balcarce, Argentina. Samples were analyzed by indirect fluorescence antibody test (IFAT) testing (27,28) and by IgG ELISA using total *T. gondii* lysate antigen (TLA). Based on these analyses, 18 sera were negative by both techniques, and 14 sera that were positive by both techniques were selected. All sera were aliquoted and stored at –20 °C until use.

### 2.5 IgG-ELISA

Sera from infected lambs were analyzed by IgG-ELISA using total *T. gondii* lysate antigen (TLA), as previously described (24). Recombinant antigens were also evaluated following established protocols (22–24). Ninety-six–well plates (Nalge Nunc International) were individually coated with 10 µg of each rAG. For the rAG combination, equimolar amounts were used: 10 µg rGra4Gra7 (45 kDa), 3 µg rGra8 (14 kDa), and 10 µg rROP2 (42 kDa). ELISAs were developed and read as described (24). Data were analyzed and graphed using GraphPad Prism version 5.0 (GraphPad Software, CA, USA).

### 2.6 Histopathology and cyst detection

Portions of the brain and spinal cord from euthanized lambs were fixed in 10% buffered formalin for 5 days, processed using standard histopathological techniques, and stained with hematoxylin and eosin. Eight tissue sections, including cervical spinal cord, medulla oblongata, pons, midbrain, thalamus, frontal and occipital cortex, and cerebellum, were examined under optical microscopy at 10×, 20×, and 40× magnifications in a double-blind manner. Lesions were scored on a scale of 0 to 3 according to the extent of pathological changes, as previously described (24): Score 0 (without lesions): no infiltration of inflammatory cells. Score 1 (mild lesions): occasional and mild infiltration of mononuclear inflammatory cells and gliosis (in some samples, forming small focal aggregations). Score 2 (moderate lesions): multifocal and moderate infiltration of mononuclear inflammatory cells and gliosis. Score 3: multifocal and severe infiltration of mononuclear inflammatory cells and gliosis accompanied by necrosis. Two independent observers evaluated each sample from each lamb, and the average score was calculated (24). These values were then summed to obtain a total injury score for each animal.

The presence of brain cysts was assessed by microscopic visualization of brain homogenates, prepared directly or after Percoll enrichment (29). Briefly, approximately 1 g of brain tissue was manually homogenized in 150–200 mL of PBS (1×) and kept overnight at 4 °C in 50 mL tubes. Supernatants were discarded, and pellets were washed in PBS and centrifuged. For Percoll enrichment, the resuspended material was layered onto 5 mL of 0.9% NaCl/90% Percoll (Sigma) and centrifuged at 1000 × g. The interface was recovered in 5 mL PBS, and 300 µL of the suspension was examined microscopically. Cysts were analyzed using the same criteria as for mouse brain samples (29). Additionally, cyst-like structures were recorded during histopathological analysis.

### 2.7 Statistical and graphics

For sera from experimentally infected lambs, optical density (OD) values obtained in IgG-ELISAs with TLA and rAGs were normalized (rOD) to the corresponding day 0 values to allow comparative analyses between rAGs and/or lamb groups. A heat map was generated using the relative rOD values of each serum in relation to the corresponding lesion score.

For sheep sera, OD values were analyzed to compare seropositive and seronegative animals. For the comparison of sera from naturally infected sheep, two-way analysis of variance (ANOVA) was performed, followed by Tukey’s multiple comparison test (p** < 0.01; p**** < 0.0001). Sensitivity and specificity were assessed using contingency tables, where Sensitivity = TP / (TP + FN) and Specificity = TN / (TN + FP). Graphs were generated using GraphPad Prism version 5.0 (GraphPad Software, CA, USA).

## 3 Results

### 3.1 Lamb Infection

To generate sera from experimentally infected animals, 3-month-old lambs were subcutaneously infected with RH and M49 *T. gondii* strains at two inoculation doses (5 × 10⁶ and 5 × 10⁷ tachyzoites). Rectal temperature was monitored during the first week post-infection (Figure 1A). In all infected groups, values exceeded 40 °C, consistent with fever induced by active *T. gondii* infection, whereas temperatures in uninfected controls remained stable at approximately 39 °C (30).

**Figure 1.**
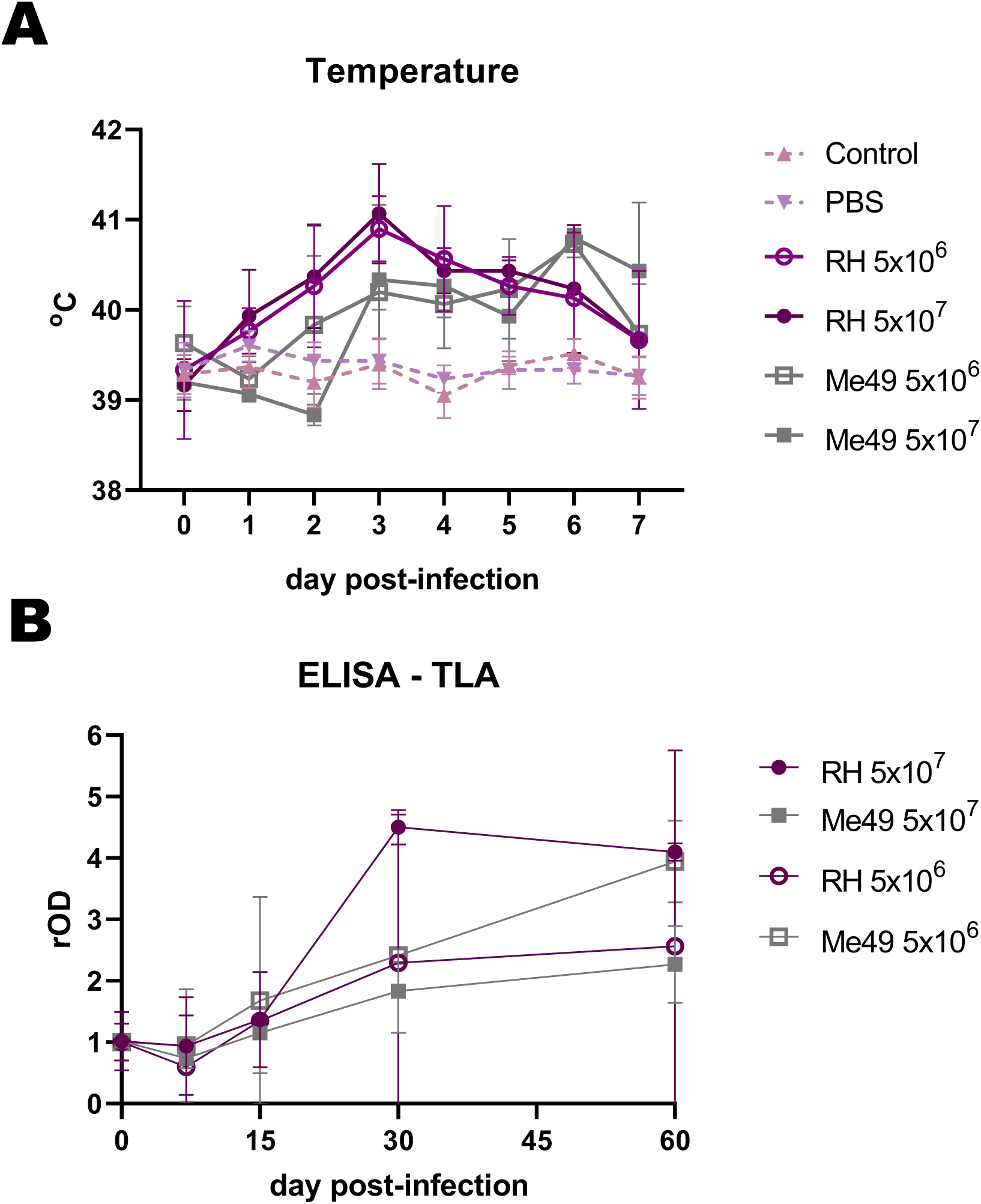
Clinical and humoral response after experimental infection of lambs with different doses (5 × 10⁶ and 5 × 10⁷) and strains (RH and Me49) of *Toxoplasma gondii*. **(A)** Rectal temperature in experimentally infected and uninfected controls. **(B)** Humoral responses were evaluated by IgG-ELISA against total lysate antigen (TLA) over the course of infection. Optical density (OD) values were normalized to day 0 readings and expressed as relative OD (rOD).

Serological responses were subsequently assessed throughout the experiment by IgG-ELISA using *T. gondii* lysate antigen (TLA). Despite some variation in mean antibody profiles between groups, a general increase in humoral response was observed from days 15–30 through day 60 post-infection (Figure 1B). One lamb from the RH 5 × 10⁶ group was excluded from all analyses due to persistently negative results, while another from the RH 5 × 10⁷ group was excluded because of a positive serum result detected at day 0.

### 3.2 Kinetics of the IgG humoral response against rAGs

The kinetics of IgG response to rGra4Gra7, rGra8, rROP2, rBCLA, and rCST9 were evaluated in the different experimental infection models, as shown in Figure 2. An early IgG response against rGra4Gra7 and rGra8 was elicited, detectable from 15 days post-infection. A chronic IgG response to rBCLA was detected from day 30 post-infection. Both IgG responses to rCST9 and rROP2 followed a similar pattern to rBCLA, suggesting their association with the chronic stage of infection.

**Figure 2.**
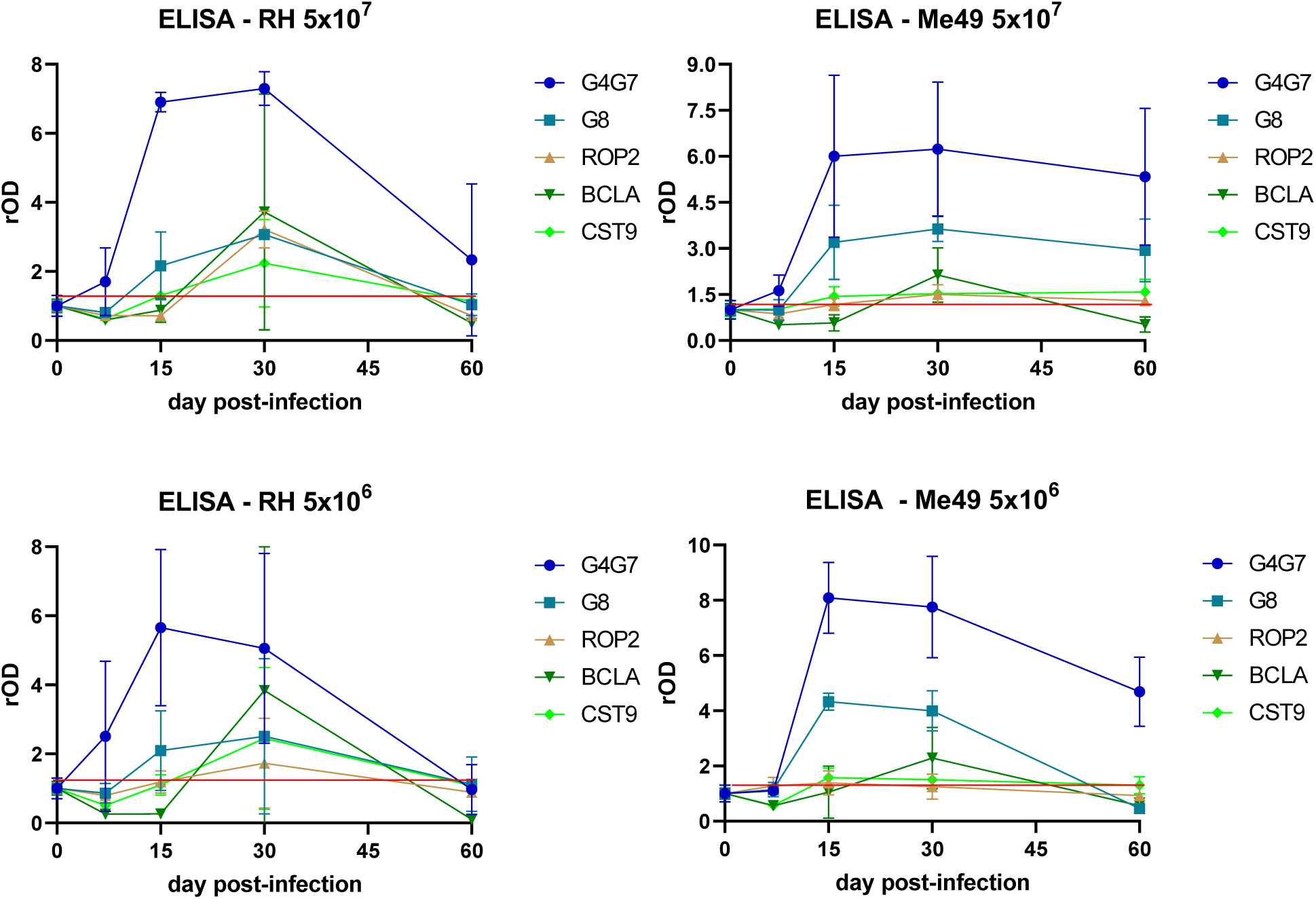
Profile of the humoral response against individual antigens. IgG reactivity to rGra4Gra7, rGra8, rROP2, rBCLA, and rCST9 was assessed by ELISA throughout the course of infection. Optical density (OD) values were normalized to the corresponding day 0 readings and expressed as relative OD (rOD). The red line represents the mean OD at day 0 plus one standard deviation.

Notably, anti-rGra4Gra7 and anti-rGra8 IgG levels declined sharply at 60 days post-infection in lambs inoculated with the RH strain, while in animals infected with Me49, antibody levels remained relatively stable throughout the experiment. Conversely, the IgG responses against rBCLA, rROP2, and rCST9 were stronger in RH-infected lambs than in those infected with Me49, although they also decreased by day 60. Infection dose had little influence on the kinetics of the response, regardless of the *T. gondii* strain used. However, a marked degree of inter-individual variability was observed among lambs, as reflected in the wide variance of antibody responses within each group (Figure 2).

### 3.3 Humoral Response and Histopathological Analysis

At 60 days post-infection, lambs were euthanized, and their brains were collected to evaluate strain- and dose-dependent differences in lesion development and pathology. *T. gondii* infection. While individual injuries were categorized from 0 (none) to 3 (severe injury), the sum of the individual averages to obtain the total injury includes not only the degree of injury but also the number of injuries observed. Therefore, it may yield values higher than 3 for a lamb (Figure 3A). Unexpectedly, no correlation was found between lesion scores and the infecting *T. gondii* strain; however, significantly higher scores were observed in animals infected with the lower inoculation doses. Cyst-like structures observed by histological analysis were predominantly observed in lambs presenting more severe lesions (Figure 3 B).

**Figure 3.**
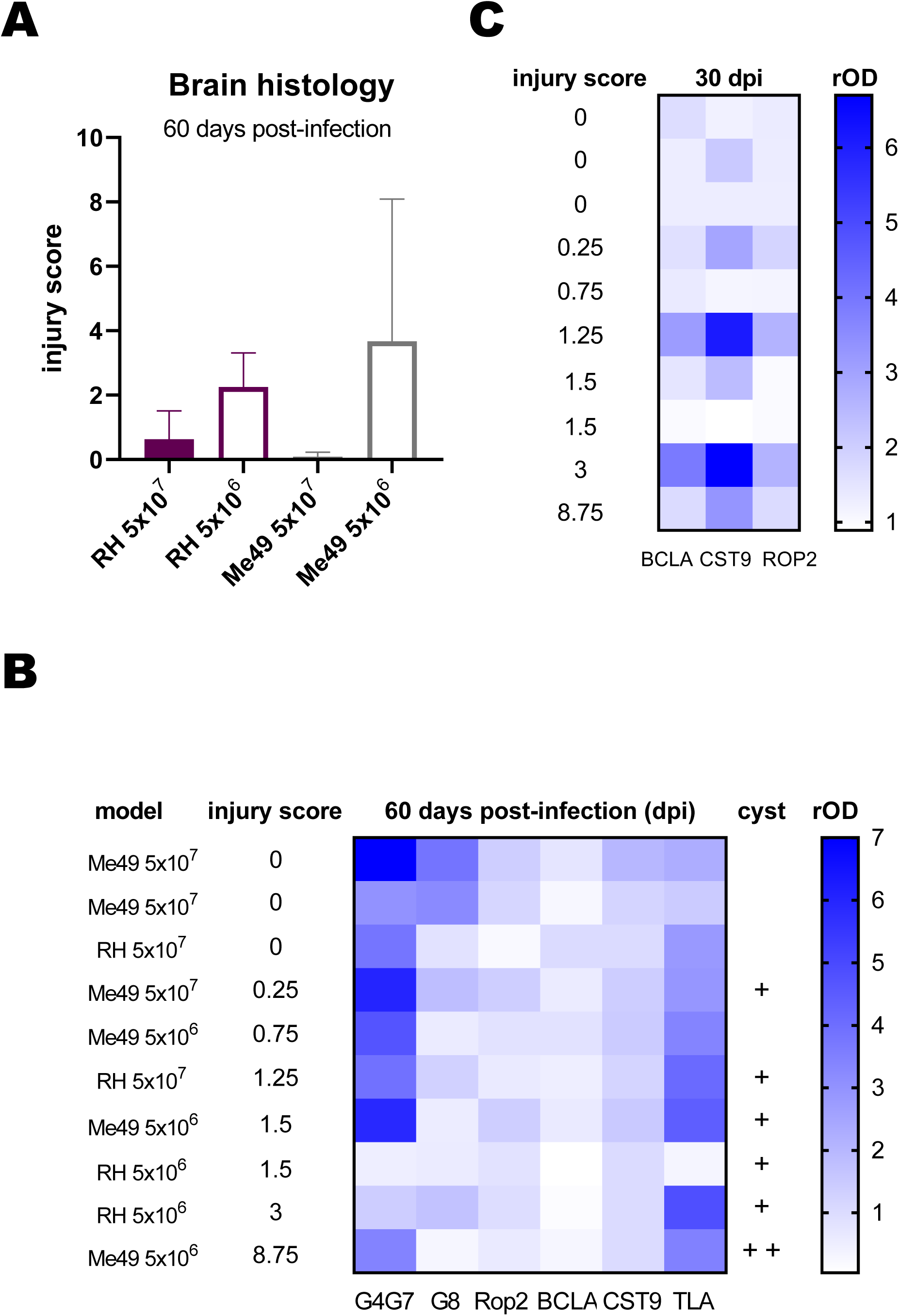
Histopathological analysis of infected lambs. Brain regions were examined by hematoxylin and eosin staining to assess lesions and inflammatory processes. Lesions were scored on a scale from 0 to 3, with 0 indicating no lesions and 3 indicating severe lesions. Scoring was performed independently by two observers. Eight brain samples were analyzed per lamb, covering comparable brain areas across all animals. **(A)** Total injury score per lamb, calculated by summing the average scores obtained by both observers for the eight samples. **(B)** Heat map of relative OD (rOD) values for each serum against the indicated rAGs at day 60 post-infection. Lambs were ranked according to their total injury score. +, indicates the presence of 1 cyst in 300 µl of brain sample or in all 8 samples of the histopathological study. **(C)** Heat map of rOD values for each serum against rBCLA, rCST9, and rROP2 at day 30 post-infection. Lambs were ranked according to their total injury score.

To explore potential associations between lesion severity and humoral responses, the IgG reactivity to rAGs was analyzed using sera collected at day 60 post-infection, and results were represented as a heat map (Figure 3B). No consistent relationship emerged across all antigens, although lambs with higher lesion scores showed a slight tendency toward lower rOD values for rGra4Gra7. To further investigate temporal dynamics, heat maps were generated for individual rAGs across multiple time points. From this analysis, IgG responses to rBCLA, rCST9, and rROP2 at day 30 post-infection displayed a clearer trend, with antibody levels paralleling the degree of tissue injury (Fig. 3C).

### 3.4 Humoral Response Against rAGs in Naturally Infected Sheep

To evaluate the diagnostic performance of rAGs in naturally infected sheep, seronegative animals and seropositive animals were analyzed. Consistent with the results obtained in experimentally *T. gondii-*infected lambs, the strongest IgG responses were directed against rGra4Gra7 (78% positive), followed by rGra8 (43%) and rRop2 (28%) in naturally *T. gondii-*infected sheep, previously tested by IFA and TLA-ELISA (*n* =18) (Figure 4A). Considerable variability was observed among individual animals, as reflected by the wide dispersion of antibody levels. When reactivity to rGra4Gra7 and rRop2 was combined, 86% of seropositive animals were detected (data not shown).

**Figure 4.**
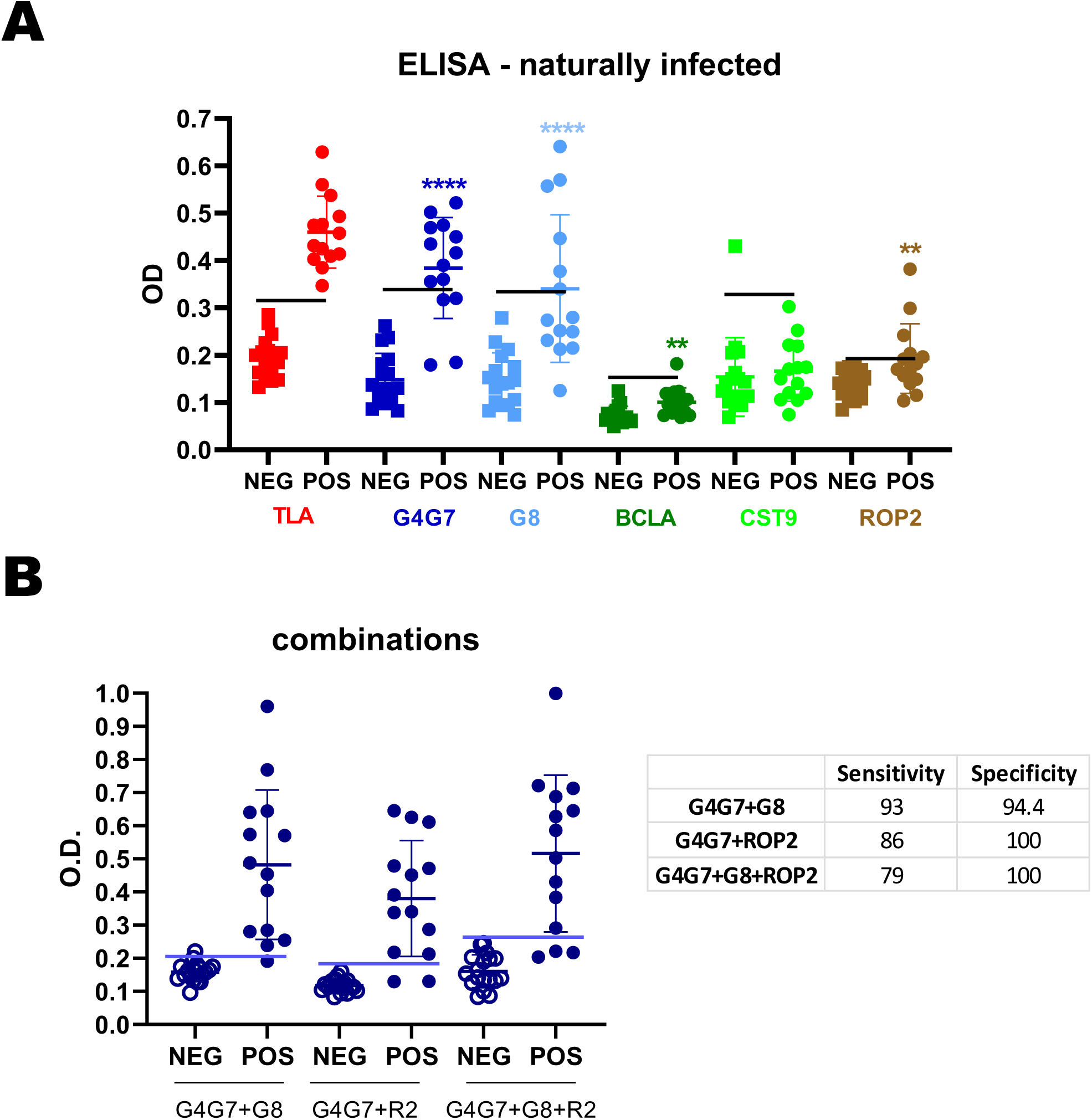
Humoral response of naturally infected sheep. **(A)** Sera from seronegative (n = 18) and seropositive (n = 14) sheep were analyzed by IgG-ELISA against individual rAGs. OD values are shown, with each point representing a single serum. Mean ± SD values for each rAG are indicated. The horizontal line denotes the cut-off for each antigen, defined as the mean OD of seronegative sera plus 3 SDs. **(B)** Combinations of rAGs were analyzed by IgG-ELISA. Sensitivity and specificity values are summarized in the table. Statistical significance: \*\***p** < 0.01; \*\*\*\***p** < 0.0001.

Different rAG combinations were tested to assess any improvement in diagnostic sensitivity (Figure 4B). Although all combinations increased overall sensitivity, comparison of sensitivity, specificity, and mean optical density values indicated that the most effective combination for detecting IgG in naturally infected sheep was rGra4Gra7 plus rGra8.

## 4 Discussion

In this study, we evaluated the humoral response to several *T. gondii* rAGs in both experimentally and naturally infected sheep. We further examined the potential association between antibody responses to the rAGs and histopathological lesions in experimentally infected lambs. This approach provides initial insights into the antigen-specific humoral response in ovine toxoplasmosis and may help identify candidate markers of infection and/or disease pathology.

The infection models employed in this study do not fully replicate the natural course of *T. gondii* infection. Under field conditions, lambs are typically infected by ingesting *T. gondii* oocysts shed into the environment by definitive hosts, with infection initiating in the intestinal epithelium. In contrast, we used a subcutaneous route of infection with *T. gondii* tachyzoites. Access to oocysts is limited for most laboratories, and the RH strain does not produce oocysts (31). While oocysts could be generated using the ME49 strain, we opted for the same tachyzoite-based model as with RH to allow for comparative analysis, given the practical restrictions on oocyst collection and handling in many laboratories. Our findings indicate that these experimental models may serve as valuable proof-of-concept systems; however, confirmation should ultimately be sought using more physiologically relevant infection models or through field studies.

Kinetic analysis of IgG responses against the rAGs revealed that rGra4Gra7 and rGra8 are elicited early during infection, consistent with previous reports identifying them as markers of the acute or active phase (22,32). In contrast, the anti-BCLA IgG response was detected from 30 days post-infection, a stage typically associated with chronic infection (25). Both rRop2 and rCST9 responses displayed kinetics similar to rBCLA, suggesting that the responses to these antigens may also be preferentially elicited during the chronic phase. For rRop2, however, there are no conclusive data linking it to a specific stage of infection. In the case of CST9, our previous analyses in mouse models and human infections indicated reactivity during the acute phase (23). Responses against CST9 is detected in both stages of the parasite life cycle, localizing to dense granules and the endoplasmic reticulum in tachyzoites, and to the cyst wall in bradyzoites (23,33–35). The divergent kinetics observed between mice, humans, and sheep suggest species-specific differences in antigen exposure to the immune system. In sheep, reactivity to CST9 was low and heterogeneous, whereas in humans it was detected in only 35% of sera from acutely infected individuals (23). These findings suggest that CST9 immunoreactivity may be influenced by host genetic or environmental factors.

Our analysis revealed that the humoral response to rAGs declined sharply by day 60 post-infection in lambs experimentally infected with the RH strain. In contrast, IgG responses against rGRA4Gra7 and rGRA8 persisted throughout the experiment in lambs infected with the Me49 strain. These findings suggest that the magnitude and duration of the IgG response to *T. gondii* in lambs are influenced by the infecting strain. They further indicate that RH and Me49 may follow distinct propagation patterns in this host: the RH strain appears to disseminate rapidly, whereas Me49 seems to spread more slowly in its transitions into a stable chronic infection.

Histopathological analysis of brain tissues and cyst detection revealed considerable heterogeneity among the lambs. Although the sample size was limited, a correlation was observed between lesion scores and the presence of brain cysts, but no differences were detected in RH and Me49-infected lambs. Unexpectedly, more severe lesions were detected in lambs that received a lower infection dose, a finding for which we currently lack a clear explanation. Additional investigations, particularly those assessing cellular and inflammatory responses, will be necessary to clarify these observations.

A limiting factor was the difficulty of detecting cysts in brain samples. In the lambs where cysts were observed, only a single cyst was found across all analyzed tissue sections, with two cysts detected in just one case. Notably, these occurrences coincided with lambs that developed strong IgG responses against rBCLA and rCST9 during the 30 days post-infection, suggesting a possible association between these antibody responses and tissue cyst burden. Although this finding was expected based on previous reports (25), longitudinal studies analyzing the kinetics of cyst formation alongside humoral responses in experimentally infected sheep are needed to clarify this relationship.

Interestingly, cysts were occasionally detected in infections with both the cystogenic Me49 strain and the non-cystogenic RH strain. It is known that RH has not completely lost its ability to form cyst-like structures either *in vitro* or *in vivo*, but rather that this capacity depends on specific conditions. For example, RH cysts have been induced in mice following treatment with atovaquone and pyrrolidine dithiocarbamate, or after passage in rats (36,37). In addition, RH tachyzoites can form cyst-like structures in vitro, either after treatment with the RAD51 inhibitor B02 or even spontaneously in skeletal muscle cell cultures (23,38). These observations make it plausible that RH can disseminate in lambs and generate some cyst-like structures. However, further studies are needed to determine whether these structures correspond to mature cysts and, importantly, whether they are capable of surviving passage through the digestive tract.

The IgG response against rGRA4GRA7 and rGRA8 appears to be associated with the acute phase of infection. At this stage, when tachyzoites are actively proliferating, most of the tissue damage caused by toxoplasmosis is expected to occur (39). Interestingly, our analysis suggested a trend in which higher levels of anti-rGRA4GRA7 IgG were associated with lower lesion scores. The role of antibodies in controlling toxoplasmosis remains poorly understood, despite evidence of their neutralizing activity against specific parasite proteins (40,41). For example, Cha et al. demonstrated that monoclonal antibodies targeting GRA6 and SAG1 reduced parasite burden in experimentally infected mice (42). Although our sample size is limited, these findings support the generation of new hypotheses. One possibility is that strong humoral responses against rGRA4 and/or rGRA7 may contribute to improved control of parasitemia and/or modulate inflammatory processes (43), thereby mitigating pathology. By contrast, higher levels of anti-BCLA and anti-CST9 IgG were more predictably associated with brain lesions linked to cyst formation (44). In mouse models, the peak of proinflammatory cytokine secretion occurs around day 30 and declines by day 60 (45). Such inflammatory responses are essential to control *T. gondii* infection (46). In the future, it will be important to assess whether humoral responses against specific rAGs, particularly rGRA4 and rGRA7, are associated not only with modulation of pathology but also with clinical outcomes such as abortion in sheep infected with *T. gondii*.

Beyond their role in shaping the immune response, these rAGs also show promise as diagnostic tools in sheep, especially when used as chimeric proteins or in combination. Similar findings have been reported by other researchers, who achieved sensitivities and specificities close to 100% in sheep and other livestock using rAG combinations (47–49). For example, an ELISA incorporating the bradyzoite antigen rBAG1 together with rGRA1, an antigen expressed during the acute phase but also localized in the cyst wall, reached a sensitivity of 95% and a specificity of 100% with sheep sera (50).

## 5 Conclusion

This study provides new insights into the humoral immune response of sheep to *T. gondii* antigens, highlighting putative pathological markers. The results indicate that specific IgG against rGra4Gra7 and rGra8 are reliable markers of acute infection, while the response to rBCLA, rRop2, and rCST9 are more associated with the chronic phase and cyst formation. Based on histological versus reactivity levels of each rAG, we consider that strong IgG responses against rGra4Gra7 may contribute to limiting pathology. Conversely, higher anti-BCLA and anti-CST9 responses correlate with tissue cyst burden and lesion severity. Beyond their immunological relevance, rGra4Gra7 and rGra8 show promise for improving diagnostic accuracy in sheep, outperforming traditional TLA-based assays. While the small sample size limits definitive conclusions, these findings generate hypotheses for further studies on immune modulation, pathology, and reproductive outcomes in ovine toxoplasmosis.

## Data availability statement

The raw data supporting the conclusions of this article will be made available by the authors, without undue reservation.

## Ethics statement

The animal study was approved by the Ethics Commission for Experimental Animal Studies of INTA CeRBAS (CICUAE 244/2022). The study was conducted in accordance with the local legislation and institutional requirements.

## Author contributions

IG, PF, LMC: Formal analysis, Investigation, Methodology, Visualization, Writing – review & editing. GJC, VS: Conceptualization, Visualization, Writing – review & editing. LMM, ES, ASC, VS, CM: Methodology, Writing – review & editing. MC, DPM, and SOA: Conceptualization, Funding acquisition, Project administration, Supervision, Writing – original draft.

## Funding

The author(s) declare that financial support was received for the research and/or publication of this article. The present work was financed by the PICT cat I-56, PIP 2021-1168, PITEs 50, INTA (PD.I116/P02), and UNMdP (AGR717/24).

## Acknowledgments

L.M. Campero, D.P. Moore, S.O. Angel, M. Clemente, and V.A. Sander are members of Consejo Nacional de Investigaciones Científicas y Técnicas (CONICET). S.O. Angel, M. Clemente, and V.A. Sander are Professors at Universidad Nacional General San Martin (UNSAM), and D.P. Moore is also a Professor at Universidad Nacional de Mar del Plata. I. Gual is a post-doctoral fellow of CONICET and a Professor at Universidad Nacional de Mar del Plata. P. Formigo is student of UNSAM. G.J. Cantón is a researcher at the National Institute of Agricultural Technology (INTA Balcarce).

## Conflict of interest

The authors declare that the research was conducted in the absence of any commercial or financial relationships that could be construed as a potential conflict of interest.

## Generative AI statement

The authors declare that no Gen AI was used in the creation of this manuscript.

## Notes

### Competing Interest Statement

The authors have declared no competing interest.

